# Noradrenergic but not dopaminergic neurons signal task state changes and predict re-engagement after a failure

**DOI:** 10.1101/686428

**Authors:** Caroline I Jahn, Chiara Varazzani, Jérôme Sallet, Mark E Walton, Sébastien Bouret

**Author notes:** Corresponding author. Team Motivation Brain and Behavior, Institut du Cerveau et de la Moelle Épinière, Hôpital Pitié-Salpêtrière, 47, boulevard de l’Hôpital, 75013 Paris, France.

## Abstract

The two catecholamines, noradrenaline and dopamine, have been shown to play comparable roles in behaviour. Both noradrenergic and dopaminergic neurons respond to salient cues predicting reward availability and to stimulus novelty, and shape action selection strategies. However, their roles in motivation have seldom been directly compared. We therefore examined the activity of noradrenergic neurons in the *locus coeruleus* and putative midbrain dopaminergic neurons in monkeys cued to perform effortful actions for rewards. The activity in both regions correlated with the likelihood of engaging with a presented option. By contrast, only noradrenaline neurons were also (i) predictive of engagement in a subsequent trial following a failure to engage and (ii) sensitive to the task state change, the discovery of the new task condition in unrepeated trials. This indicates that while dopamine is primarily important for the promotion of actions directed towards currently available rewards, noradrenergic neurons play a crucial complementary role in mobilizing resources to promote future engagement.

## Introduction

Catecholaminergic neuromodulation is thought to be critical for numerous aspects of behaviour, including motivation, learning, decision-making and behavioural flexibility (Robbins & Roberts 2007; Doya 2008; Sara 2009; Robbins & Arnsten 2009; Sara & Bouret 2012). Both noradrenaline and dopamine neurons respond to novel and salient stimuli and signal predictions of future reward (Schultz 1998; Bouret & Sara 2004; Ravel & Richmond 2006; Berridge 2007; Ventura et al. 2007; Matsumoto & Hikosaka 2009; Bromberg-Martin et al. 2010) and both systems have been implicated in motivating action (Robbins & Everitt 2007; Nicola 2010; Bouret et al. 2012; Varazzani et al. 2015; Jahn et al, 2018; Walton & Bouret, 2019). Nonetheless, the specific contributions of dopamine and noradrenaline to these functions remain unclear, in part as their roles have seldom been compared in the same task (but see Bouret et al. 2012 and Varazzani et al. 2015).

*Locus coeruleus* (LC) noradrenergic-containing neurons have a long-stated role in signalling new information about the state of the world, specifically a change in predictability of the environment (Swick et al, 1994; Vankov et al, 1995; Dalley et al, 2001; Aston-Jones & Cohen, 2005; Bouret & Sara, 2005; Yu & Dayan, 2005). LC neurons are particularly sensitive to unexpected and/or novel stimuli (Kety 1972; Foote et al. 1980; Aston-Jones & Bloom 1981; Grant et al, 1988; Sara & Segal, 1991; Vankov et al, 1995; Bouret & Sara, 2004; Bouret et al, 2012), and the transient activation of LC neurons in response to unexpected stimuli is often thought to facilitate adaptation through an increase in behavioural flexibility (Bouret & Sara, 2005; Dayan & Yu, 2006, Einhauser et al, 2008; Nassar et al, 2012, Urai et al. 2017; Muller et al. 2019). In that frame, the magnitude of LC responses to sensory stimuli increases when these stimuli are unexpected, and therefore provide information about the state of the world that may be useful to guide subsequent behaviour. By contrast, perfectly expected stimuli provide little information, and so their presentation should not require the updating of behaviour. In other words, such a function could allow the activation of LC neurons to promote the adaptation of behaviour in response to a change in the state of the world (Aston-Jones & Cohen, 2005; Bouret & Sara, 2005; Yu & Dayan, 2005). Such a role for noradrenaline in behavioural flexibility has received strong support from pharmacological studies (Devauges & Sara, 1990; Tait et al, 2007; McGaughy et al, 2008; Jahn et al, 2018; Jepma et al, 2018).

More recently, noradrenaline function has been extended to include the promotion of effortful actions (Ventura et al. 2008; Bouret & Richmond 2009; Zénon et al. 2014; Varazzani et al. 2015). Indeed, LC neurons are reliably activated when animals initiate an action (Bouret & Sara, 2004; Rajkowski et al, 2004; Kalwani et al 2014). Critically, the magnitude of this activation seems to be related to the amount of effort necessary to trigger the action (Bouret & Richmond, 2015; Varazzani et al, 2015). In line with this interpretation, we recently used a pharmacological manipulation to demonstrate directly that, on top of its role in behavioural flexibility, noradrenaline was also causally involved in motivation (Jahn et al, 2018). One interpretation of the dual role of noradrenergic LC neurons in behavioural flexibility and motivation is that flexibility relies upon their response to unexpected stimuli whereas their role in motivation relies upon their activation at the triggering of effortful actions. Alternatively, the response of LC neurons to unexpected stimuli could be directly related to motivation.

Since the tripartite relationship among LC activity, processing of expected vs unexpected stimuli, and motivation remain unexplored, we re-analysed a data set of noradrenergic neurons in the LC recorded in monkeys presented with cues signalling how much effort they would need to expend to gain rewards of various sizes (Varazzani et al. 2015). The task was designed such that rejecting an offer caused it to be re-presented on the subsequent trial, and the analyses reported by Varazzani et al. (2015) deliberately excluded such repeated trials. Here, by including those trials, we could investigate separately (i) the sensitivity to task state changes in unrepeated vs. repeated trials and (ii) the encoding of motivational processes, by examining the modulation of LC activity by willingness to perform the presented option (engagement) in the current or in the future trials.

Moreover, to gain further insight on the specific role of noradrenaline as compared to dopamine neurons, we compared the activity of LC neurons to that of putative DA neurons recorded from *substantia nigra pars compacta* and *ventral tegmental area* (SNc/VTA) in the same paradigm. Indeed, dopamine is also implicated in novelty and information seeking (Horvitz et al. 1997; Schultz 1998; Costa et al. 2014; Bromberg-Martin & Hikosaka, 2009; Naudé et al. 2016), as well as playing a prominent role in motivation and action initiation (Walton & Bouret, 2019). As for LC noradrenergic neurons, we could examine separately the relation between dopaminergic neurons and sensitivity to task state changes and willingness to perform the presented option.

We found that that although the magnitude of the neuronal response at the cue predicted the engagement in effortful actions similarly in the two catecholaminergic systems, only noradrenaline neurons were sensitive to changes in task state, i.e. to the difference between repeated (and therefore perfectly expected) and unrepeated (and therefore informative) stimuli. Moreover, while dopamine neurons only reflected the engagement at the cue onset, noradrenaline cells were also activated by erroneous fixation breaks, in a manner that predicted the likelihood of future engagement after erroneous trials. Taken together, our analyses demonstrate complementary but distinct roles for noradrenaline and dopamine in signalling new states of the world and in motivating current or future engagement with effortful actions.

## Results

### Behaviour

Three monkeys were trained to perform a task in which visual cues indicated the amount of effort (3 effort levels) that was required to obtain a reward (3 reward levels) (fig 1A and B). Effort and reward levels were manipulated independently across the 9 task conditions. On a given trial, monkeys could either engage in the effortful action (whether action is correct or not) or fail to engage by breaking fixation (the proportion of trials where monkeys maintained fixation and omitted the response was negligible). Importantly, unsuccessful trials, which effectively represent a failure, were repeated (see Material and Methods and figure 1 for details).

**Figure 1:**
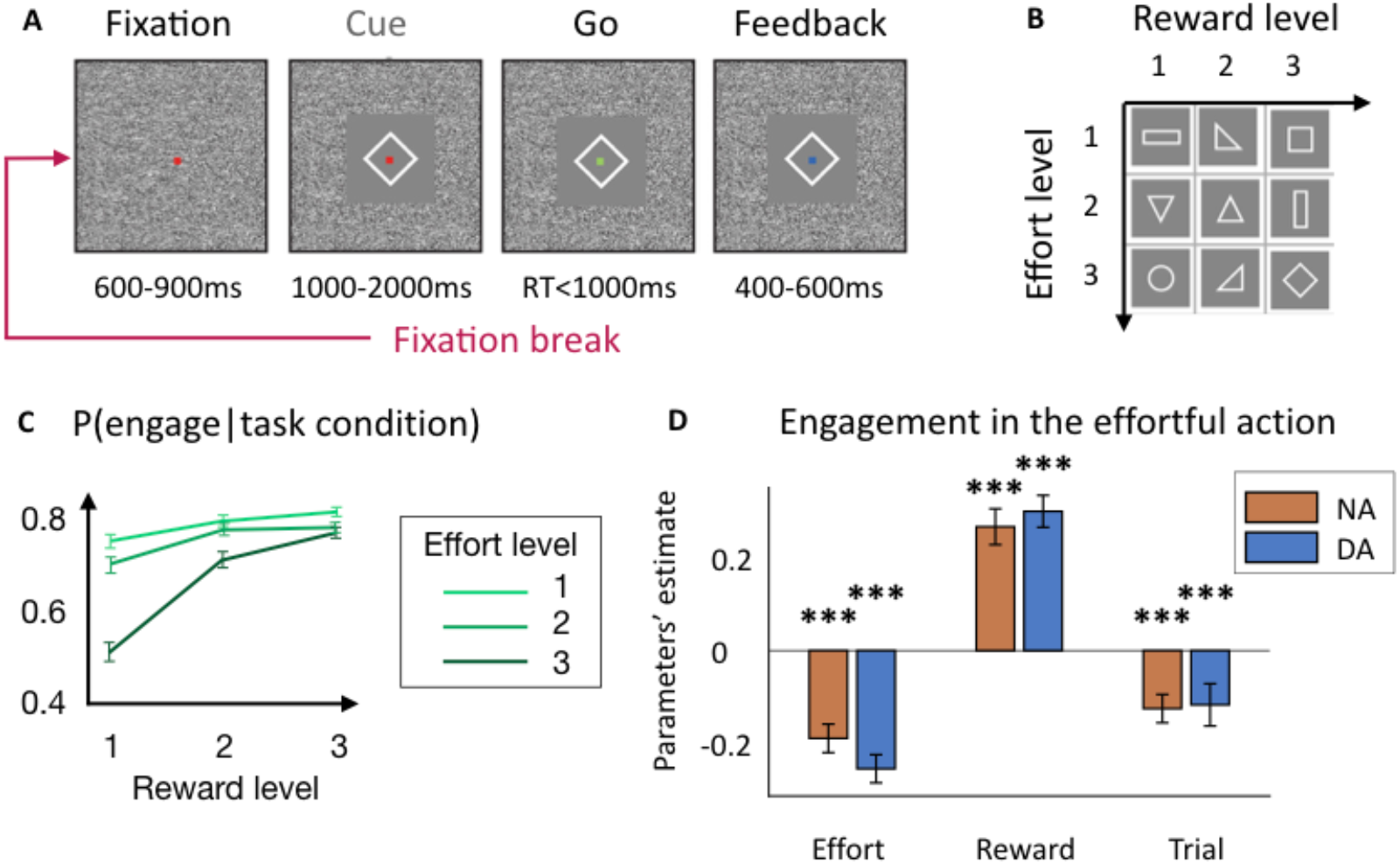
Task and behaviour. A) Task structure. Monkeys had to squeeze a clamp with a certain minimum intensity to obtain reward of a certain magnitude. During the whole trial, monkeys had to maintain fixation on a dot at the centre of the screen. If they broke the fixation, the trial restarted from the start after an inter-trial interval delay. A trial started with monkeys fixating the red dot, then a cue appeared indicating the effort and reward levels for the current trial. The dot turned green (Go signal) and monkeys had to squeeze the clamp to the minimum force threshold indicated by the cue. Upon reaching this threshold, the dot turned blue (Feedback) and remained blue as long as monkeys had to keep on squeezing. If monkeys maintain the effort long enough, they received the amount of reward indicated by the cue. B) Task design. Each trial corresponded to one of nine experimental conditions, defined by three levels of effort and three levels of reward. C) Probability to engage with the action as a function of effort and reward levels. Computed for all NA and DA sessions together. D) Weights of the task parameters in the decision to engage with the effortful action. Multi-level logistic regression of the decision to initiate the action by the three experimental task parameters: effort level, reward level and trial number. Significant negative effect of effort level (p<0.001) and trial number (p<0.001) and significant positive effect of reward level (p<0.001) in both NA and DA session (no difference between NA and DA sessions for all three parameters (p<0.05)). *** p ≤ 0.001.

The monkeys’ willingness to engage in the task – measured as the attempt to squeeze the clamp after seeing the cue – was clearly affected by the information about the upcoming effort and reward levels (task condition) of the trial (fig 1C-D). In both sessions when noradrenergic (NA) or dopaminergic (DA) neurons were recorded from, the likelihood of engagement in the effortful action was negatively affected by the effort level (NA: β=-0.19±0.03, t(91)=-6.19, p<0.001; DA: β=-0.26±0.03, t(83)=-8.43, p<0.001) and positively modulated by the reward level (NA: β=0.27±0.04, t(91)=6.93, p<0.001; DA: β=0.31±0.04, t(83)=8.78, p<0.001). Moreover, monkeys’ engagement was negatively modulated by the trial number (NA: β=-0.13±0.03, t(91)=-4.11, p<0.001; DA: β=-0.12±0.05, t(83)=-2.58, p<0.001) (fig 1D). Note that there was no significant difference between effort level, reward level and trial number weights in engagement across for NA and DA recording sessions (p=0.13, p=0.52 and p=0.88 respectively). This was confirmed by a 2-way ANOVA measuring the effect of task factor (effort and reward) and recording type (NA or DA) onto –β(effort) and β(reward): main effect of task factor F(1,348)=3.35, p=0.07) but no main effect of recording session type (F(1,348)=2.14, p=0.15) and no interaction (F(2,348)=0.23, p=0.63), meaning that engagement was affected in the same way by the two task factors in both types of recordings.

### Noradrenergic and dopaminergic neurons’ activity reflects monkeys’ engagement in the task

We have seen previously that the task factors (i.e. effort level, reward level and trial number) influenced the probability of monkeys to engage with the effortful action. Therefore, we first measured the influence of these task factors on neurons’ activity at the time of cue. Dopaminergic neurons’ activity was significantly positively modulated by reward level (β=0.05±0.01, t(83)=3.67, p<0.001) and negatively modulated by the effort level (β=-0.02±0.001, t(83)=-2.01, p=0.05), as well as by trial number (β=-0.06±0.03, t(83)=-2.53, p=0.01) (fig. 2A). Noradrenergic neurons’ activity was only significantly modulated by the reward size (β=0.04±0.001, t(91)=4.05, p<0.001) but not reliably modulated by either the effort level (β=-0.01±0.01, t(91)=-1.15, p=0.25) nor trial number (β=-0.03±0.03, t(91)=-1.02, p=0.31) (fig 2A). However, we found no significant difference between the encoding of the effort level and the trial number between dopaminergic and noradrenergic neurons (p=0.42 and p=0.37 respectively). Critically, there was a significant difference between the weights of effort and reward in the firing rates of both noradrenergic and dopaminergic neurons (2-way ANOVA measuring the effect of task factor (effort and reward) and recording type (NA or DA) onto –β(effort) and β(reward): main effect of task factor F(1,348)=9.71, p=0.02) but no main effect of recording session type (F(1,348)=0.61, p=0.4) and no interaction (F(2,348)=0.04, p=0.8). This means that the relative sensitivity of noradrenergic and dopaminergic neurons to the task factors was similar, with a greater sensitivity for reward than effort (post-hoc T-test on the distribution of –β(effort) and β(reward): t(350)=-3.13, p=0.002).

**Figure 2:**
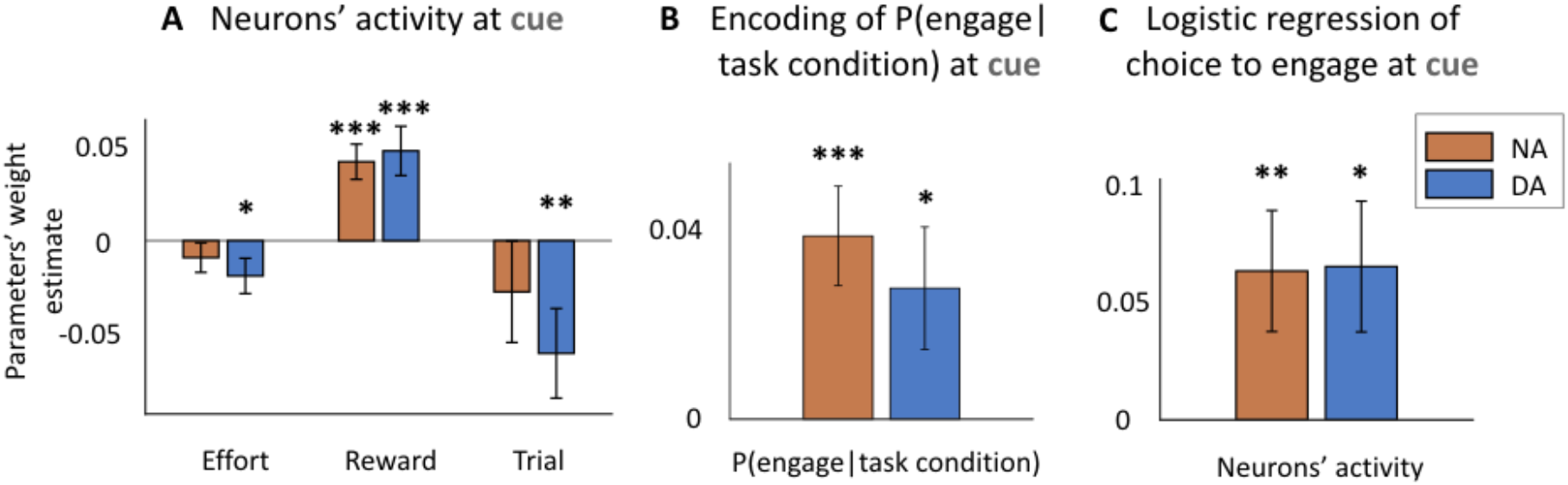
noradrenergic and dopaminergic neurons encoding of the task parameters and engagement at the time of cue. A) Encoding of task parameters at the time of cue (0-500ms from cue onset). Dopaminergic neurons were sensitive to all three task parameters (effort level: p=0.05; reward level: p<0.001; trial number: p=0.01). Noradrenergic neurons were only significantly sensitive to the reward level (p<0.001). No significant difference between the encoding of effort level and trial number in noradrenergic and dopaminergic neurons (p>0.05). * p < 0.05; ** p ≤ 0.01; *** p ≤ 0.001. B) Noradrenergic and dopaminergic neurons reflect the engagement in a task condition. Linear regression of the probability to engage in a given task condition (effort and reward levels) for each session. Both populations encode significantly the probability to engage (p<0.05), no difference between the strength of encoding across populations (p>0.05). * p < 0.05; ** p ≤ 0.01; *** p ≤ 0.001. C) Noradrenergic and dopaminergic neurons’ activity reflects the engagement on a trial-by-trial basis throughout the session. Logistic regression of Noradrenergic and dopaminergic neurons’ activity on engagement in the action. Both populations predict the engagement in the action (p<0.05). * p < 0.05; ** p ≤ 0.01; *** p ≤ 0.001.

After having considered the relation between neuronal activity and task factors, we looked at the relationship between neuronal activity and the engagement in the effortful action. First, we did it across the nine task conditions (defined by a combination of effort and reward levels) by using an aggregate measure of the engagement for each condition (the probability to engage given the task condition). This tested whether neuronal activity directly reflected the probability for the monkeys to engage in a particular task condition. For each recording, we regressed this z-scored probability of engagement on neurons’ activity and found a significant positive effect at the population level, for both noradrenergic and dopaminergic neurons (NA: β=0.04±0.01, t(91)=3.70, p<0.001, DA: β=0.03±0.01, t(83)=2.16, p=0.03) (fig 2B). Again, there was no difference in the strength of this signal encoding between populations (p=0.50). Moreover, this activity was specific to the onset of the cue as there was no significant encoding of this probability before the cue onset (pre-cue period) even in repeated trials, in which monkeys already knew which cue was coming (500ms window before cue onset: NA: p=0.17, DA: p=0.71). We also examined the relation between neuronal activity and engagement on a trial by trial basis. We found that both noradrenergic and dopaminergic responses were predictive of engagement on a trial by trial basis (NA: β=0.06±0.03, t(91)=2.47, p=0.01; DA: β=0.06±0.03, t(83)=2.36, p=0.02) (fig 2C). Here again, there was no difference in the strength of this signal encoding between dopaminergic and noradrenergic neurons (p=0.96). Moreover, the activity was specific to the onset of the cue, with no encoding of engagement in the pre-cue period (NA: p=0.08, DA: p=0.88).

Overall, we found that even if, contrary to behaviour, the activity of the noradrenergic and dopaminergic systems is biased toward the encoding of reward compared to effort the firing of these neurons reflected the engagement in the effortful action in a similar fashion at the time of the cue.

### Both noradrenergic and dopaminergic neurons encode monkeys’ engagement, but only noradrenergic neurons are sensitive to changes in task state

In order to understand if catecholaminergic neurons also encode changes in task states (i.e. when their responses to cues differed between repeated and non-repeated trials) and to determine the relationship between this factor and motivation (engagement), we compared the encoding of these two variables at the time of cue. To examine the effect of changes in task states, we compared cue-evoked activity in repeated (‘non-informative cue’) versus non-repeated (‘informative cue’) trials. Since erroneous trials were repeated, and monkeys knew the structure of the task, they could predict following an error that the same condition (with the same visual cue) would be presented again, such that the visual cue provided no information about the task state. By contrast, after a correct trial, any of the nine task conditions could be pseudo-randomly presented to the monkey, such that visual cues now provided information about the upcoming reward and effort levels (task state). Erroneous trials were mainly of two types: (i) monkeys broke the fixation (no engagement) and (ii) monkeys engaged (tried to squeeze the clamp) but did not execute the action correctly. Therefore, as not all trials in which monkeys engaged were successful, we were able to look conjointly at the effect of engagement and the information being presented on neuronal activity.

First, we found no interaction between the linear encoding of the effort, reward levels and trial number with whether the trial was repeated or not in either noradrenergic neurons or dopaminergic neurons (see Materials and Methods, NA: p=0.24, p=0.26 and p=0.58 respectively; DA: p=0.26, p=0.27 and p=0.10 respectively). This means that the task condition was encoded in a similar fashion whether the cue was informative or not.

To examine the effect of engagement and task state change above and beyond the effect of a particular task condition (effort and reward levels), we regressed out the effect of the task condition on the firing rate of neurons and looked at the effect of engagement and task state change (unrepeated vs. repeated trials) on the remaining neuronal activity (see Material and Methods). Here, we found an important dissociation between the activity of noradrenergic and dopaminergic neurons (fig 3). For a given trial condition, noradrenergic neurons were more active either when the action was initiated (vs not) *or* when the cue provided information about the new task condition (in unrepeated vs repeated trials) in a given experimental condition (β(engagement)=0.11±0.03, t(91)=3.40, p<0.001; β(task state change)=0.16±0.04, t(91)=4.23, p<0.001). We also found a significant negative interaction (β(interaction)=-0.06±0.02, t(91)=-3.02, p=0.003), which indicates that engagement and information effects were not perfectly additive: when both factors were combined, the firing rate increased less than by the sum of the two separate effects. On the other hand, while dopaminergic neurons were on average more active when monkeys engaged in a given condition (β=0.08±0.04, t(83)=2.05, p=0.04), they were *not* sensitive to the task state change (p=0.56). There was also no significant interaction between the two effects (p=0.36), and the main effects were similar when we removed the interaction. A direct comparison of these effects between noradrenergic and dopaminergic neurons confirmed that, while there was no difference in the strength of their encoding of engagement in the task (p=0.59) noradrenergic neurons encoded significantly more task state change than dopaminergic neurons (p<0.001).

**Figure 3:**
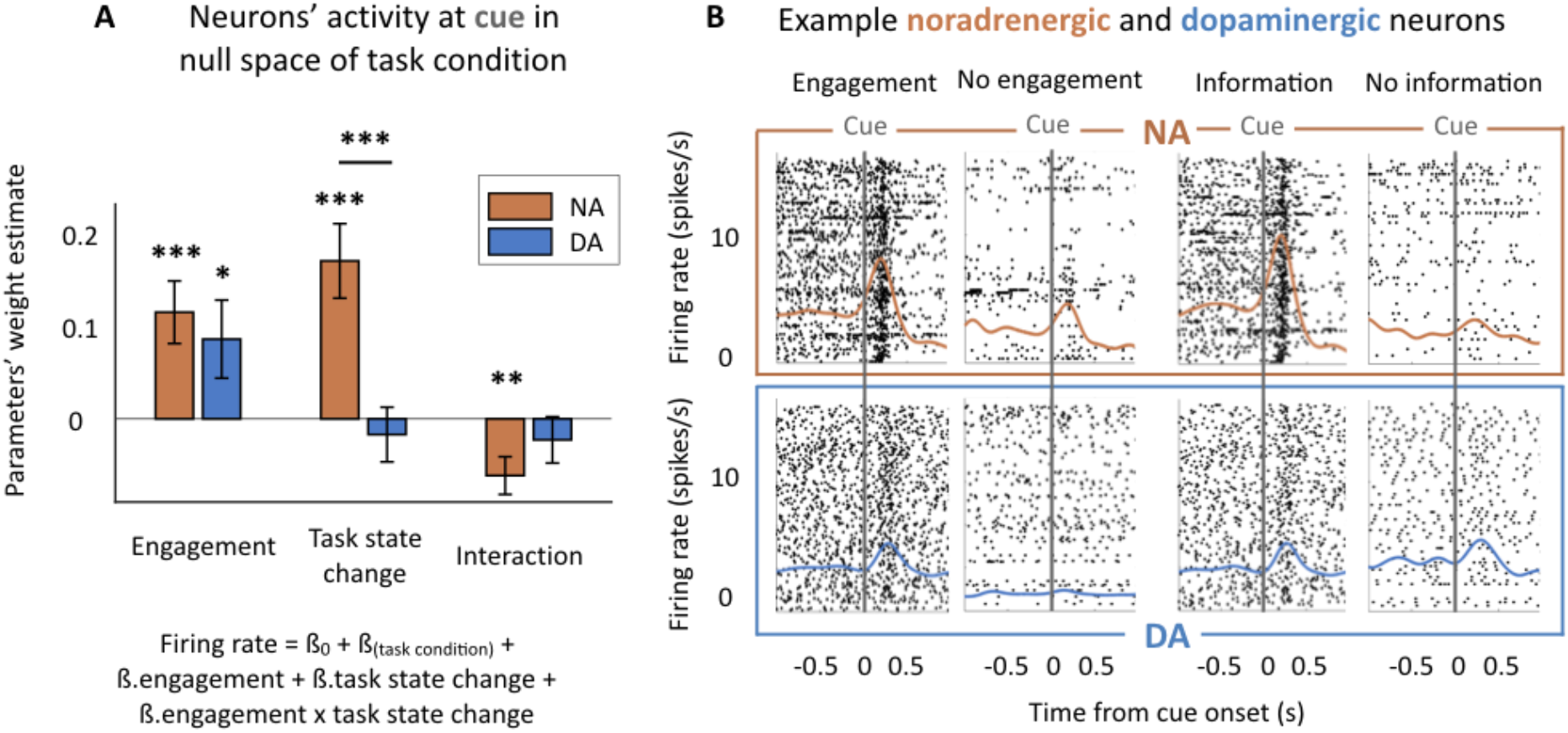
Change in task condition is encoded by noradrenergic but not dopaminergic neurons. A) Encoding of engagement and trial repetition in null space of task condition at cue (0-500ms from cue onset). Noradrenergic neurons encoded significantly the change in trial condition, the engagement and the interaction (all p<0.01). Dopaminergic neurons encoded only significantly the engagement (p<0.05). * p < 0.05; ** p ≤ 0.01; *** p ≤ 0.001 B) Example noradrenergic and dopaminergic neurons. Neuronal activity of two representative neurons around the cue onset (grey vertical line). Top: spike activity (raster and spike density function) of a noradrenergic neuron showing a strong activation at cue. The activation is stronger in engaged vs. non-engaged trials (all experimental conditions pooled together) and for informative vs. non-informative cues. Bottom: same but for a dopaminergic neuron showing an intermediate activation at cue onset. Its activity was greater for engaged than non-engaged trials but was not modulated by the task state change of the cue. Note, even though the baseline firing appears different in these example neurons, there was no reliable effect of engagement before cue onset. Each panel corresponds to a different number of trials (each trial is a line in the raster plot).

Here again, this effect was specific of the onset of the cue as when we examined the 500ms pre-cue period, there was neither an effect of engagement (NA: p=0.17, DA: p=0.77) nor an effect of task state change (NA: p=0.96, DA: p=0.07). There was also no effect of engagement in the pre-cue period if we only examined repeated trials where monkeys already knew the task condition (NA: p=0.31, DA: p=0.47). In short, when comparing the encoding of engagement and task state change (unrepeated vs. repeated trials) variables over and above the task variables, both noradrenergic and dopaminergic neurons encoded the engagement in the task, but only noradrenergic neurons encoded the task state change (whether the cue was informative or not). In addition, these effects were unaffected by the addition of trial number to the analyses, which captures the influence of fatigue and satiety (main effects of engagement and task state change remained as described before; main effect of trial number: NA: p=0.47, DA: p=0.02; interaction of engagement and task state change with trial number did not reach significance in either noradrenergic or dopaminergic neurons, NA: p(engagement)=0.84, p(task state change)=0.97, DA: p(engagement)=0.91, p(task state change)=0.19) (see supplemental figure 1A). Thus, engagement and task state change had specific effects on neurons’ firing rates, which in turn were independent of the progression in the session.

### Only noradrenergic neurons were activated after a failure to engage and are sensitive to the task condition

We next examined the activity of dopaminergic and noradrenergic neurons time-locked to fixation break, which resulted in trial abortion. We focused our analysis on three epochs: a baseline epoch from −600 to −300ms prior to fixation; a pre-fixation break epoch corresponding to the 300ms prior to fixation break, and post fixation break epoch corresponding to the 300ms following fixation break. There was neither a significant activation of dopaminergic neurons before fixation break (p=0.62) nor after the fixation break (p=0.49). By contrast, noradrenergic neurons were significantly activated after (mean difference=0.30±0.09 spikes/s, t(83)=3.31, p=0.001), but not before (p=0.81) the fixation break had occurred. This activation corresponds to an average change of 16.5%±0.04 of activity between before (average firing rate = 2.83 spikes/s) and after (average firing rate = 3.12 spikes/s) the fixation break (fig 4A). At the single neuron level, 18.1% noradrenergic neurons were activated at the fixation break (one-tailed T-test: firing rate(pre fixation break) < firing rate(post fixation break), p<0.05 were considered as significant). Note that all results hold true if we removed fixation break events that occurred less than 500ms after the cue onset.

**Figure 4:**
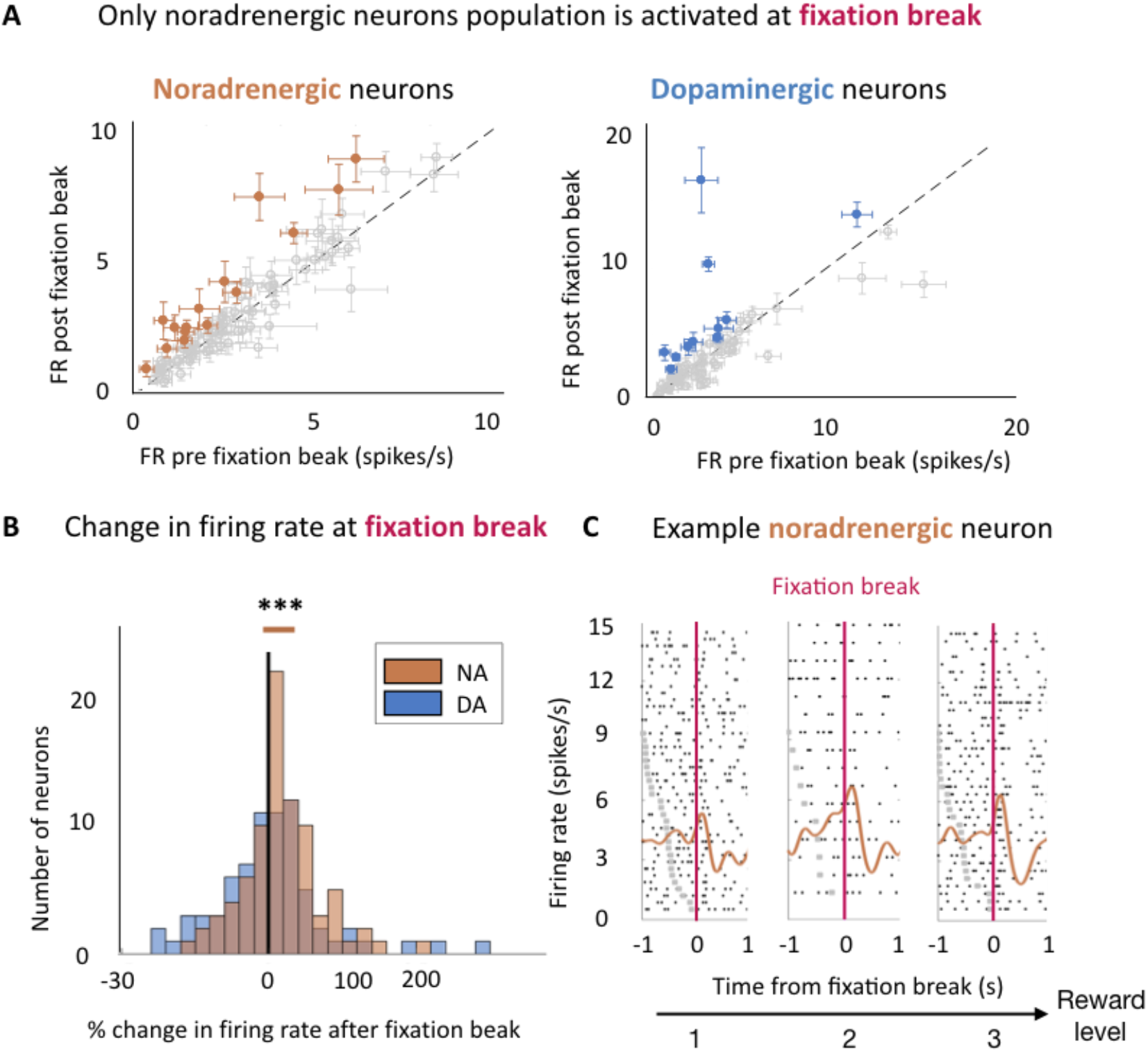
Noradrenergic but not dopaminergic neurons were activated after the fixation break. A) Only noradrenergic neurons population is activated at fixation break. Firing rate pre (−300 – 0ms) and post (0 – 300s) fixation break for both noradrenergic (left) and dopaminergic neurons (right). Points and error bars are mean ± SEM. Solid points indicate a significant activation (One-tailed T-test, p<0.05). For illustration purposes only, we have removed two dopaminergic neurons (with a non-significant activation at fixation break), whose firing rates were above 20 spikes/s from the display. B) Noradrenergic and Dopaminergic neurons’ change in firing rate evoked by activity after fixation break (0-300ms from fixation break). The distributions are represented on a log-scale. Noradrenergic neurons population was significantly activated after the fixation break (p=0.001) but not dopaminergic neurons population (p=0.49). *** p ≤ 0.001. C) Example noradrenergic neurons at fixation break for each reward level. Neuronal activity representative of noradrenergic neuron around fixation break (pink vertical line). Trials are sorted by decreasing latency between cue onset (grey dots) and fixation break. Cue onset is only visible for bottom trials, with latencies shorter than the displayed 1 sec. Spike activity (raster and spike density function) of a noradrenergic neuron showing an increase after the fixation break. In addition, its activity is modulated by the reward level (p<0.001).

We then looked at the modulation of fixation-break related activity across task conditions. The firing of dopaminergic neurons did not show any significant modulation across task conditions (probability to engage with the task condition: p=0.97) or behavioural responses (engagement in the next trial: p=0.45) and it will not be described further. By contrast, noradrenergic neurons’ evoked activity was positively modulated by the reward size (β=0.06±0.02, t(83)=3.64, p<0.001) but neither by the effort level nor by the trial number (β(effort level)=-0.01±0.02, t(83)=-0.91, p=0.37; β(trial number)=-0.04±0.03, t(83)=-1.31, p=0.20) (fig 4B). Note however, that the difference between the sensitivity to effort and reward did not reach significance (t-test on –β(effort) and β(reward): t(166)=1.88, p=0.06). This activity was specific to the onset of the fixation break as there was no modulation of the activity by these task factors in the 300ms before the fixation break (effort level: p=0.50; reward level: p=0.15; trial number: p=0.9).

### Noradrenergic neurons activity predicted the engagement on the next trial

Finally, we examined the relationship between fixation-break evoked activity and the probability, across sessions, that the monkeys engaged on the next trial. Here again, we only looked at fixation break events that occurred after cue onset, meaning that the monkeys always knew the task condition at the time of the fixation break.

We found a significant positive effect of the probability to engage given the task condition on LC activity at the time of the fixation break (β=0.05±0.02, t(83)=2.79, p=0.007). In other words, the more monkeys tended to engage in a specific task condition, the more noradrenergic neurons would be active if a fixation break occurred in this task condition. This effect was also present in the pre-fixation break activity (−300-0ms to fixation break) (β=0.15±0.06, t(83)=2.55, p=0.01), suggesting that it appeared after cue onset, in line with the fact that noradrenergic neurons also displayed a positive relation with task engagement at the time of the cue onset (fig 2B). Indeed, we found a significant positive correlation (r=0.33, p=0.002) between the strength of the encoding of the probability to engage at the time of cue and at the time of the post-fixation break (fig 5B). In short, noradrenergic neurons were activated both at cue onset and at the fixation break when it occurred. They tended to be more active in conditions associated with a greater probability of engagement, both at the cue onset and at the time of fixation break, and these two responses were correlated across the population of LC neurons.

**Figure 5:**
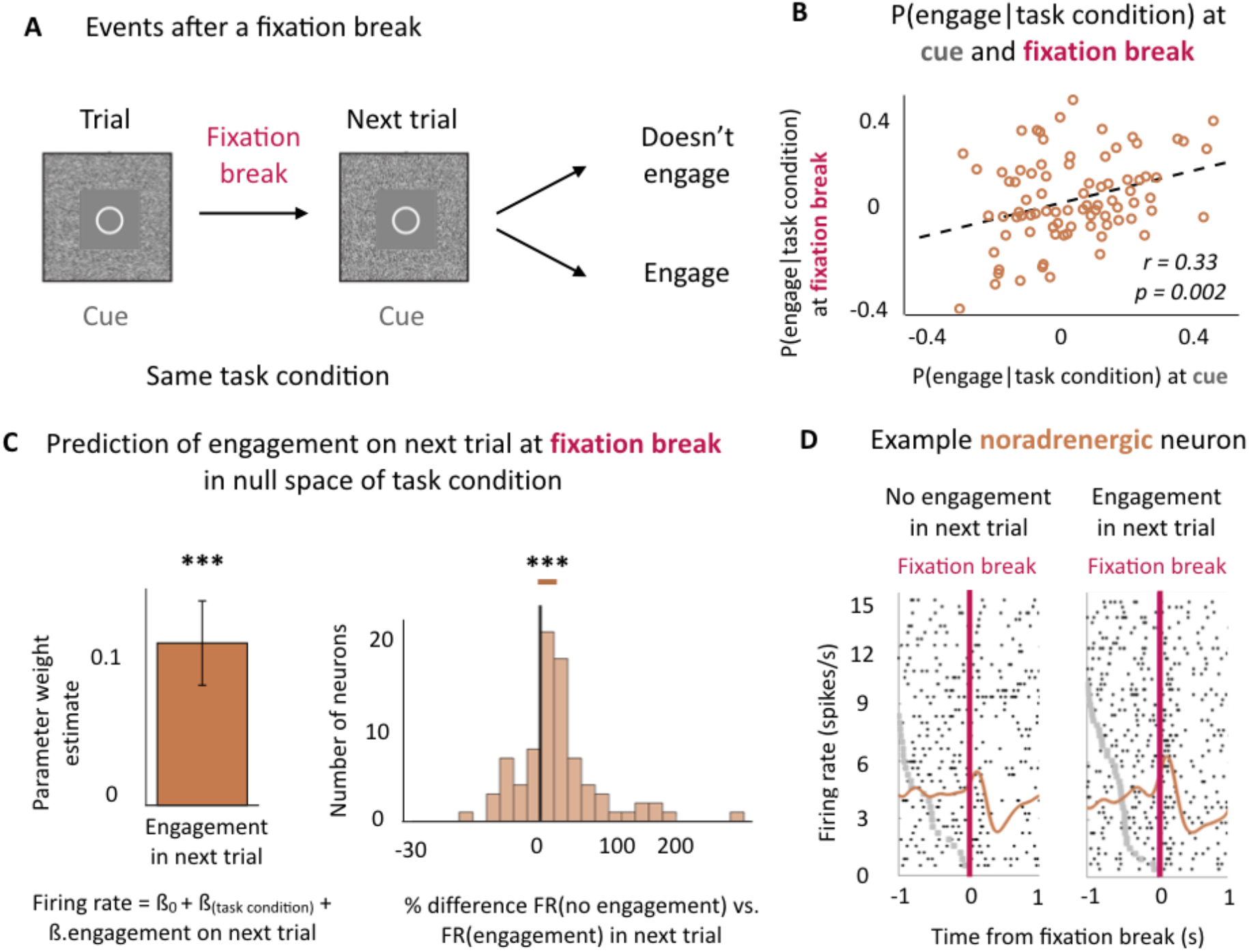
Noradrenergic neurons’ activity predicts the engagement on the next trial. A) Task structure after a fixation break. B) Correlation between noradrenergic neurons’ encoding of the probability to engage for each task condition at the cue onset and the fixation break across sessions. Significant correlation (r=0.33, p<0.01). C) Noradrenergic neurons’ activity at fixation break is predictive of engagement in the next trial above and beyond the task condition. Linear regression, significant effect (p<0.001). % difference between the firing rate distribution for no engagement in next trial and engagement in next trial in the null space of task conditions (mixed effect linear regressions on non-z-scored distributions). The distribution is represented on a log-scale. Significant difference (p<0.001). *** p ≤ 0.001. D) Example noradrenergic neurons at fixation break for no engagement (left) and engagement (right) in the next trial. Neuronal activity (raster and spike density function) is displayed around fixation break (t=0, pink vertical line). Trials are sorted by decreasing latency between cue onset (grey dots) and fixation break. Cue onset is only visible for bottom trials, with latencies shorter than the displayed 1 sec. As a majority of LC neurons, this one shows a stronger activation when monkeys engaged on the next trial (p<0.001).

Given this strong relation between LC activity and probability of engagement in the current trial when monkeys erroneously break fixation, we were interested to examine whether this activity could also predict monkeys’ likelihood of engagement in the following trial. After a fixation break, two things could happen on the next trial (and therefore in the same task condition): monkeys could now choose to engage with the same task condition or could again reject the offer (fig 5A). We therefore examined if LC activity at the time of fixation break could provide information about engagement in the next trial, over and above task condition.

In fact, the magnitude of the fixation-break activation of noradrenergic neurons (controlled for task condition) was predictive of subsequent engagement in the next trial (β=0.12±0.003, t(83)=3.84, p<0.001; effect calculated on the z-scored distributions of firing rates and translating to an average difference of 25.1%±0.1 of activity between non-engage and engage on the next trial conditions) (fig 5C). At the single neuron level, only 6.5% of neurons showed a significant effect (compared to 7.6% of neurons showing a significant sensitivity to reward at fixation break and 20.6% at cue). Hence, although the effects seen at the fixation break are relatively weak at the single neuron level, they are very consistent across the population, such that at the population level the effect clearly reaches significance. In fact 66.3% of neurons showed small but consistently greater activation in trials in which monkeys engage on the next trial, which is comparable to the proportion of neuron displaying a positive relation with reward at the fixation break (63%) or at the cue (66.3%). We controlled for potential interactions with confounding factors such as task state change (whether the erroneous trial was itself a repeated or not), trial number and their interactions with the effect of the engagement in the next trial, but none of them were significant (main effects: p(information)=0.18, p(trial number)=0.15; interactions with engagement with next trial: p(information)=0.27, p(trial number)=0.81). As previously mentioned, this activity was specific of noradrenergic neurons as dopaminergic neurons were not activated post-fixation break and did not signal the engagement in the next trial (p=0.45) (see supplemental figure 1B).

Finally, we looked whether the effect of the engagement in the next trial could be found before the cue of the next trials. In other words, we looked if we could predict the engagement before the cue (−500 – 0ms) for trials where a fixation break occurred. We found that it was not the case (p=0.25) and could therefore only conclude that noradrenergic neurons predict the engagement on a trial-by-trial basis.

In summary, we found that noradrenergic but not dopaminergic neurons’ activity at fixation break reflected the probability to engage both in the current and in the subsequent trial, over and above cost-benefit task conditions.

## Discussion

In this task, monkeys were presented with informative (non-repeated) and non-informative (repeated) cues instructing them to produce actions of different intensities to gain rewards of different magnitudes. The probability that monkeys would try to produce the action (engagement) depended on the task condition (effort and reward levels) but failing to engage would only lead to the repetition of the same task condition. Repeated trials constituted series of actions towards the same goal: the reward. This goal directed behaviour ended when the goal was reached. From that perspective, there is a clear transition in behaviour after a correct trial, as animals get started on another trial, another goal directed behaviour (Bouret & Richmond 2009). Hence, given the structure of the task, unrepeated trials are more likely to constitute a task state changes than repeated ones from a goal-directed behaviour perspective. We used this task structure to reveal the precise roles of noradrenergic and dopaminergic neurons in encoding motivation to engage in the task and in signalling task state changes. We used the engagement in a task condition on a specific trial as a measure of motivation and found that both noradrenergic and dopaminergic neurons’ activities were predictive of the engagement. Their activities were not only correlated with the session-average probability to engage in a particular task condition, but also with the trial-by-trial engagement. Furthermore, their activities were correlated with engagement over and above the specific task condition. This strengthens the role of both catecholaminergic systems in motivating effortful, reward directed actions.

However, the activity of noradrenergic and dopaminergic neurons differed significantly when it came to signalling task state changes. First, only noradrenergic neurons’ activity was sensitive to whether or not the visual cue was providing information about the new task state (which was the case only in non-repeated trials), over and above its relation with upcoming reward and effort levels. Moreover, noradrenergic, but not dopaminergic, neurons displayed activity after a fixation break, which ended the trial and represented a failure to engage. This activity scaled with the probability of engagement given the task condition and it was positively correlated with the engagement in the next trial. Hence, noradrenaline, contrary to dopamine, plays a role both in signalling information about task state and in promoting current and future effortful actions given this information.

### Similarities and dissimilarities of the role of the catecholaminergic systems in motivation

This study builds on experiments presented in Varazzani et al (2015), but here includes both repeated and non-repeated, and correct and incorrect trials, rather than just the non-repeated correct trials reported in Varazzani et al (2015). This allowed us to examine the influence of information about task state changes and motivation to engage, and not just the cost-benefit parameters of the presented cues, on neural activity. The inclusion of these additional trials did lead to slight differences in the strength of encoding of task parameters to those reported previously. However, importantly the overall pattern of effects was comparable, and any differences were negligible compared to the difference in terms of sensitivity in noradrenaline and dopamine neurons to changes in task state.

Both noradrenergic and dopaminergic neurons’ activity was related to the engagement in the effortful actions. Dopaminergic neurons’ activity was tightly linked with the engagement in the rewarded course of action independently of whether the trial was repeated or not. Dopaminergic neurons were also activated at the time of producing the action, but contrary to noradrenergic neurons, they did not correlate with the actual force produced (Varazzani et al. 2015). The causal role of dopamine in incentive processes has been shown in different species, with an emphasis on its role in controlling reward sensitivity (Denk et al. 2005; Hoskins et al. 2014; Le Bouc et al. 2016; Yohn et al. 2016; Zénon et al. 2016; Noritake et al, 2018). Moreover, our results are in line with studies demonstrating that dopamine release is strongly driven by the initiation of a purposeful action for reward (Phillips et al. 2003; Roitman et al., 2004; Syed et al. 2016).

Noradrenergic neurons’ activity was also linked to the engagement in the effortful course of action as well as to the actual production of the action (Varazzani et al., 2015). This is in line with previous demonstrations that LC neurons respond to stimuli predicting future rewards and action initiation responses (Bouret & Sara, 2004; Bouret & Richmond 2009, 2015; Kalwani et al. 2014). Contrary to dopamine, causal manipulation of the noradrenergic system does not seem to affect incentive processes (Hoskins et al. 2014; Jahn et al. 2018). Indeed, our recent study showed that the noradrenergic system controls the amount of force produced during the action, but not the selection nor the initiation of the action (Jahn et al. 2018). Hence, the noradrenergic system might be critical to ensure that the effortful action is appropriately performed once a decision to engage has been taken (Bouret & Richmond 2015; Varazzani et al. 2015), whereas dopamine is instead key for signalling the subjective future reward to be gained by performing an action and promoting that response (Ishiwari et al., 2004; Gan et al. 2010; Pasquereau & Turner 2013; Varazzani et al. 2015; Papageorgiou et al., 2016; Salamone et al. 2016).

### Why are dopaminergic neurons not sensitive to the information about task state change in our task?

Dopamine neurons have long been reported to respond to salient novel stimuli (Strecker & Jacobs 1985; Ljunberg et al. 1992; Horvitz et al. 1997; Menegas et al. 2017) and to be implicated in novelty seeking (Costa et al. 2014). Therefore, it may initially seem surprising that in our task, dopaminergic neurons were not sensitive to the novelty of the presented task condition information. However, there are a number of important differences between these experiments and the current one. For instance, in previous experiments examining novelty seeking, it is unclear whether dopaminergic neurons are encoding new information based on the change in uncertainty about the world, independent of choice, or as a variable driving the behaviour. While Bromberg-Martin and Hikosaka showed that dopaminergic neurons were sensitive to the advanced information about the size of the reward, importantly in their study, monkeys showed a preference for obtaining this information, implying that it was therefore relevant for guiding the behaviour (Bromberg-Martin & Hikosaka 2009; Charpentier et al. 2018). In another experiment, Naudé and colleagues showed that mice preferred a probabilistic outcome to a deterministic outcome, and that this preference was controlled by the dopaminergic system (Naudé et al. 2016). These two studies show that dopaminergic neurons are sensitive to information as a variable that can influence choices through preferences, since it acted as a reward (Charpentier et al. 2018). In our task, as the cost-benefit cues were all well known, information (as provided by the cues in non-repeated, but not in repeated trials) would neither cause sensory surprise (as cues themselves were not novel) nor be relevant for modulating future choices. Therefore, although we cannot rule out that some individual dopamine neurons do code for this factor, it seems that dopamine neurons as a population do not encode the information about task state changes when this is not relevant to guide the behaviour.

### Noradrenergic neurons’ activity reflects the role of noradrenaline in information processing and engagement after a failure

The crucial difference between dopaminergic and noradrenergic neurons was that noradrenergic neurons were sensitive to the repetition of a trial at cue. Because task state changes only occur after a successful trial, lower activation of LC neurons at cue on repeated trials could reflect the fact that an error just occurred. However, we found no significant effect of error on the previous trial in baseline activity before the cue. Therefore, it is unlikely that there is a carry-over effect of error on the next trial. This lower activation in repeated trials could also be simply due to the repetition of a visual cue. However, there was no significant difference in the sensitivity to the task factors (effort and reward levels) in repeated and non-repeated trials. Hence, there is no evidence in our data for a simple stimulus repetition suppression effect. Moreover, from a goal directed behavior perspective, there is much more likely to be a state transition after a sequence ended with a reward, which would argue against a simple cue repetition response. Therefore, we attributed this lower activation to the fact that the monkeys already knew the task condition in repeated trials. Noradrenergic neurons would be sensitive to the information about task state changes, which corresponds to the discovery of a new state of the world either at the time of cue (i.e., which task condition has been selected for the current trial) but also at fixation break (an error means that the trial is terminated and that the same task condition is coming next).This is in line with the long-stated, if underspecified, role of noradrenaline in signalling important events in the environment (Kety 1972; Foote et al. 1980; Aston-Jones & Bloom 1981; Abercrombie & Jacobs 1987; Berridge & Waterhouse 2003; Vazey et al. 2018). Noradrenaline has been implicated in signalling a need to provoke or facilitate a cognitive shift to adapt to the environment (Bouret & Sara 2005; Yu & Dayan 2005; Glennon et al. 2019). Here, noradrenergic neurons’ sensitivity to change in task state at the time of cue could reflect a need to process the information about the current task condition.

Crucially, only noradrenergic neurons were activated following a break in fixation, which represents a failure to engage in the effortful action. Similar patterns of activity at the break of fixation have also been observed in mid-cingulate cortex (MCC), here modulated by how close to reward delivery the error occurred or how much effort was already invested in the task (Amiez et al. 2005). Given the connections between LC and MCC, this suggests that MCC and LC might well interact when required to signal salient events. A break of fixation was an important event not only as it signalled the end of the trial, but also the re-occurrence of same task condition in the next one. This post-fixation break activity was tightly linked to firing rates at the time of cue, which in turn reflected the probability of engagement in the effortful action. A potential scenario is that if the activity at the cue was too small to enable maintenance of the fixation and the engagement in the trial, then activity at the fixation break reflects a prospective update to enable performance of the action on the subsequent trial. Indeed, we found that when we controlled for task condition, noradrenaline neurons were more active after fixation break when monkeys then engaged in the subsequent trial. Finally, as we were never able to predict the engagement in the trial from the baseline activity at the cue, even for repeated trials and even for trials following a fixation break, we only conclude that noradrenergic neurons predict the engagement on a trial-by-trial basis but have no evidence that they do so through a slow fluctuation of activity that lasts beyond the range of a trial.

Together, these results are compatible with the idea that noradrenergic neurons signal and potentially facilitate the need to engage resources to undertake and complete effortful actions (Bouret et al. 2012; Walton & Bouret 2019). In both cases, they do it as a function of new information about the state of the world: about the start of a new and unpredictable experimental condition that will bring a reward at the cue, and about the failure to complete a trial that might has been worth it, since they re-engage immediately at fixation break.

To conclude, our data show the specific and complementary roles of dopamine and noradrenaline in motivation and behavioural flexibility. The former would promote actions directed towards currently available rewards, while the latter could play a critical role in facing challenging situations by mobilizing resources based on new information about the environment.

## Materials and Methods

### Monkeys

Three male rhesus monkeys (Monkey D, 11 kg, 5 years old; Monkey E, 7.5 kg, 4 years old; Monkey A, 10 kg, 4 years old) were used as subjects for the experiments. During testing days (Monday to Friday), they received all their water as reward on testing days and they received water according to their physiological needs on non-testing days. All experimental procedures were designed in association with the Institut du Cerveau et de la Moelle Epiniere (ICM) veterinarians, approved by the Regional Ethical Committee for Animal Experiment (CREEA IDF no. 3) and performed in compliance with the European Community Council Directives (86/609/EEC).

### Task

The behavioural paradigm has previously been described in detail in Varazzani et al. (2015). In brief, each monkey sat in a primate chair positioned in front of a monitor on which visual stimuli were displayed. A pneumatic grip (M2E Unimecanique, Paris, France) was mounted on the chair at the level of the monkey’s hands. Water rewards were delivered from a tube positioned between the monkey’s lips. Behavioural paradigm was controlled using the REX system (NIH, MD, USA) and Presentation software (Neurobehavioral systems, Inc, CA, USA).

The task consisted of squeezing the grip to a minimum imposed force threshold to obtain rewards, delivered at the end of each successful squeeze (fig 1A and B). At the beginning of each trial, subject had to fixate a red dot at the centre of the screen before a cue appeared. The cue indicated the minimum amount of force to produce to obtain the reward (3 force levels) and the amount of reward at stake (3 reward levels: 1, 2 and 4 drops of water). After a variable delay (1500±500ms from cue display), the dot at the centre of the cue turned green (Go signal) and subject had 1000ms to initiate the action, meaning squeezing the clamp very little (threshold set to detect any attempt to perform the action). If the monkey reached the minimum force threshold indicated by the cue, the dot tuned blue and remained blue if the effort was sustained for 500±100ms. At the end of this period, if at least the minimum required effort had been maintained, the water reward was delivered.

Fixation of the central dot had to be maintained through the different phases of the task. A trial was incorrect if: (i) the monkey broke fixation before the reward delivery, (ii) he squeezed the clamp before the go signal, (iii) he failed to squeeze the clamp at all or (iv) at the minimum force threshold or (v) didn’t maintain the effort long enough. After an error the same trial was repeated until it was successfully completed. Within a session, the nine combinations of effort and reward conditions were selected with equal probability and presented in a random order. As erroneous trials were repeated, the policy with the highest reward rate was to always engage until satiety.

Monkeys were trained for several months on this task. They first learned to distinguish and perform two different force levels and the difficulty of the task was progressively increased until they were could do so with the nine experimental conditions. Finally, they learned that they had to fixate the central dot to go through a trial.

### Electrophysiological recordings

Single unit recording using vertically movable single electrodes was carried out using conventional techniques. The electrophysiological signals were acquired, amplified (x10,000), digitized, and band-pass filtered (100 Hz to 2 kHz) using the OmniPlex system (Plexon). Precise description of the recording procedures can be found in the article where LC and SNc/VTA data used here were originally reported (Varazzani et al. 2015). Noradrenergic neurons recordings were performed on monkey A (29 neurons in 15 sessions) and monkey D (63 neurons in 38 sessions), midbrain dopaminergic neurons recordings were performed on monkey D (56 neurons in 38 sessions, sometimes simultaneously as noradrenergic neurons recordings) and monkey E (28 neurons in 19 sessions).

### Data analysis

Data were analysed with Matlab software (MathWorks). Figures represent data ± standard deviation to the mean.

In all our analyses we only considered trials (correct and incorrect) in which monkeys did not break the fixation before the onset of the cue (NA: 324 trials on average for monkey A and 281 for monkey D, DA: 314 trials on average for monkey D and 274 for monkey E). We took all those trials and computed the probability that for a given effort and reward level (or a given task condition), subjects would engage with the trial. We considered that monkeys engaged if they maintained fixation throughout the trial and initiated the action even if it occurred before the Go signal, (5% of trials in both noradrenergic (NA) and dopaminergic (DA) neurons recording sessions), not strongly (0% and 0.1% of trials in NA and DA sessions respectively) or long enough (8% and 10% of trials in NA and DA sessions respectively). Although it was possible to fail to engage with a trial by maintaining fixation but not squeezing the clamp, this type of mistake was rare (2% and 1% of trials in NA and DA sessions respectively) and monkeys mostly rejected a trial by breaking fixation (20% of all trials in both NA and DA sessions). Erroneous trials were therefore mainly of two types: i) monkeys broke the fixation and failed to engage with the trial (*no engagement* and *no new information* as the same trial type is presented again: 20% of all trials in both NA and DA sessions) and ii) monkeys engaged (tried to squeeze the clamp) but did not complete the correct action (*engagement* but *no new information*: 17% and 20% of engaged trials, which corresponds to 13% and 15% of all trials in NA and DA sessions respectively).

We examined the effects of effort, reward and trial number on the engagement in the action using a multi-level logistic regression for each session. The three variables were z-scored so that we could compare their weights across sessions. We then went on to examine task conditions influenced neuronal activity. To assess the effect of task conditions on neurons’ activity at the time of cue onset, we used a window from 0 to 500ms from cue onset. When we looked at these effects in the pre-cue period, we used a window from −500 to 0ms from cue onset. Neurons’ activity was measured in firing rates (spikes per second) and were z-scored scored for each session to compare the activity across neurons. First, the effects of the task factors: effort, reward and trial number in a session on neurons’ activity were estimated using a multi-level linear regression for each neuron. Second, we assessed the relationship between neurons’ firing rates and engagement in a given trial by running a logistic regression of neurons’ firing rates on engagement. Finally, we looked at the linear encoding of the z-scored probability to engage given the task condition on neurons’ firing rates using a linear regression.

When we looked at the effect of the novelty of the trial state (here referred to as “task state change”) on neuronal activity, we first looked at whether the fact that a cue was informative (I=1) or not (I=0) changed the sensitivity of neurons for the task factor (E, R, N) at the time cue by regressing the task factors and the interaction between the task factors and the informativity (I=0 or 1) onto the trial-by-trial neurons’ activity. A significant interaction would mean that an informative cue (signalling the new task state) would increase of decrease the sensitivity for the task factor. We then wanted to assess the conjoint effect of engagement and task state change on neurons’ firing rates above and beyond the effect of effort and reward levels. To do so, we ran a multi-level linear regression taking into account the task condition variability. In other words, we removed from neurons’ firing rates the effect of the task condition using a mixed model:

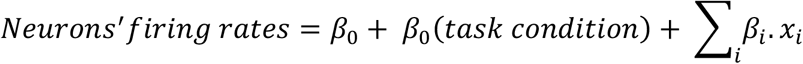

where *β_0_* a constant, *β_0_(task condition)* a constant fitted for each combination of effort and reward level (9 possibilities), *x_i_* the experimental factors and *β_i_* their weights in the linear regression (e.g. engagement, task state change, interaction). When looking at the effect of engagement and task state change at cue, we tested the following experimental factors: engagement, task state change and interaction between effect. We then added to the regression the following confounds: trial number, interaction between trial number and engagement and interaction between trial number and task state change. All results hold when adding the confounds.

We then moved on to assess whether noradrenergic and dopaminergic neurons were activated before the fixation break. We only considered fixation breaks that occurred after the display of the cue. We compared firing rates from 600ms before the fixation break to 300ms after (in 300ms windows). For all analyses at fixation break, we only included sessions during which there were more than 20 fixation break events after the onset of the cue (91 % of NA sessions and 89 % of DA sessions). Delays between the onset of the cues and fixation break events followed a Poisson-like distribution of median 845ms for NA session and 713ms for DA sessions (statistically different, t-test on the mean of the log-transformed distributions: p<0.001). To ensure that the activity at the fixation break was not contaminated by the cue response, we also looked only at fixation break events that occurred at least 500ms after the cue onset (83 % of NA sessions and 75 % of DA sessions). However, all main results were similar both with and without exclusion of the early fixation break events. To assess whether neurons were activated at the fixation break, we compared the difference in firing rate before and after the fixation break and the % of change in firing rate (by dividing by the firing rate before the fixation break). We ran a similar analysis to assess whether neurons were activated before the fixation break. When looking at the modulation of the evoked activity a fixation break, we used the same methodological approach as for the analysis of activity at cue onset. When looking at the effect of engagement in the next trial at fixation break cue, we tested the following experimental factors: engagement in the next trial. We then added to the regression the following confounds: task state change (in the current trial), trial number, interaction between the effect of engagement in the next trial and task state change and interaction between the effect of engagement in the next trial and trial number. All results hold when adding the confounds. To assess the size of the effect of engagement in the next trial, we ran the linear regression of the effect of engaging in the next trial while taking into account the task condition on the non-z-scored firing rate of neurons at fixation break and divided the regression coefficient (difference between engage and non-engage conditions) by the fixed intercept (mean firing rate across both conditions).

Second-level analyses were performed by comparing the distributions of regression coefficients against zero or other distributions (paired t-test and unpaired t-test respectively or ANOVA). Statistical reports include means of the distribution ± standard deviation to the mean, t-values or F-values and p-values.

## Acknowledgments

This research was funded by the ERC BIOMOTIV, the Paris Descartes University doctoral and mobility grants and the Wellcome Trust fellowships (MEW: 202831/Z/16/Z, JS: 105651/Z/14/Z). The Wellcome Centre for Integrative Neuroimaging is supported by core funding from the Wellcome Trust (203139/Z/16/Z).

## Supplemental figure

**Supplemental figure 1:**
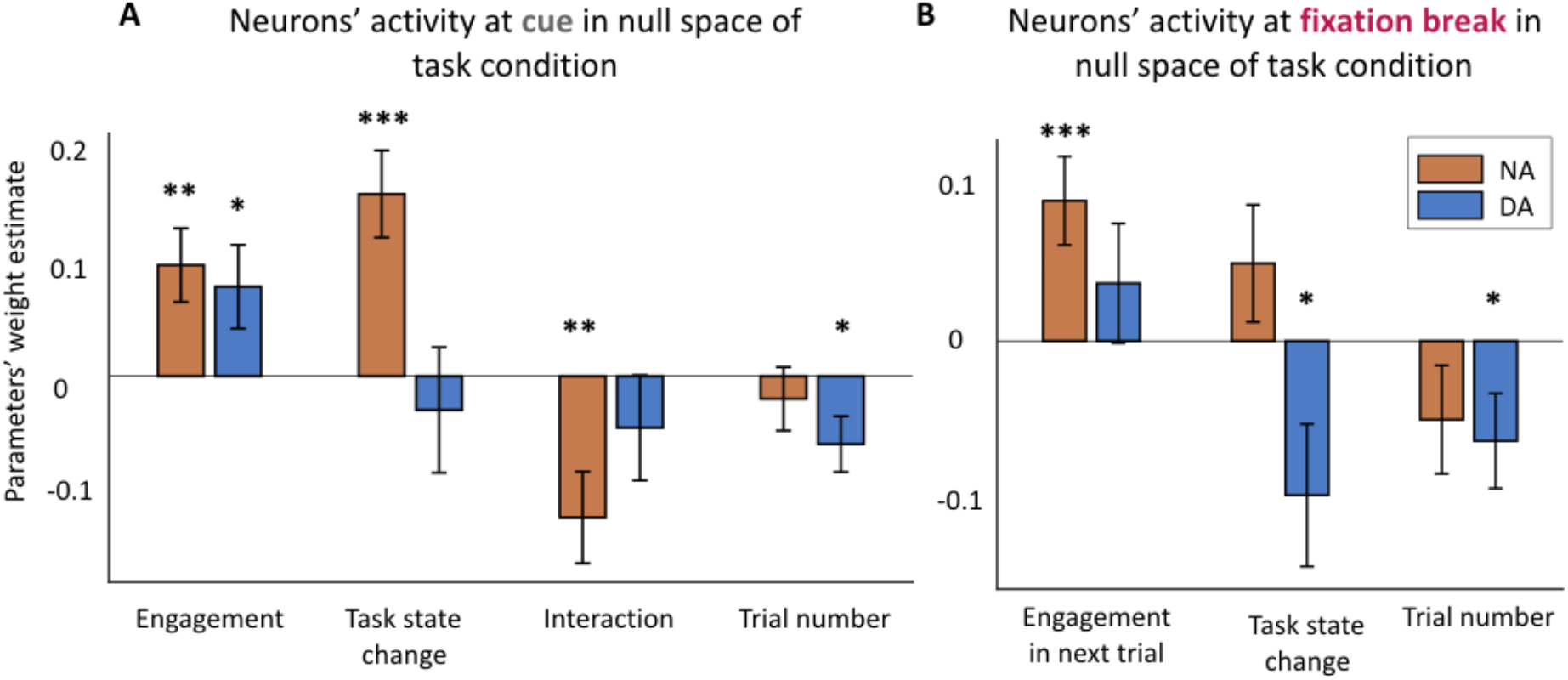
Confounds do not affect the effects described at cue and fixation break. A) Encoding of engagement, task state change and trial number in null space of task condition at cue (0-500ms from cue onset). Noradrenergic neurons encoded significantly the task state change, the engagement and the interaction (all p<0.01). Dopaminergic neurons encoded only significantly the engagement (p<0.05) and the trial number (p<0.05). Interactions between trial number and engagement and trial number and task state change were non-significant for both populations (all p>0.19). * p < 0.05; ** p ≤ 0.01; *** p ≤ 0.001 B) Encoding of engagement in the next trial in null space of task condition at fixation break (0-300ms from fixation break). Noradrenergic neurons encoded significantly the engagement in the next trial (p<0.001) even when we added the confounds: task state change and trial number (both p<0.15). Dopaminergic neurons were not significantly activated at the fixation break. However, their activity was negatively modulated by the task state change (p=0.04) and the trial number (p=0.04). Interactions between trial number and engagement in next trial and task state change and engagement in next trial were non-significant for both populations (all p>0.23). * p < 0.05; ** p ≤ 0.01; *** p ≤ 0.001

